# Evolution of corticosteroid specificity for human, chicken, alligator and frog glucocorticoid receptors

**DOI:** 10.1101/036665

**Authors:** Yoshinao Katsu, Satomi Kohno, Kaori Oka, Michael E. Baker

## Abstract

We investigated the evolution of the response of human, chicken, alligator and frog glucocorticoid receptors (GRs) to dexamethasone, cortisol, corticosterone, 11-deoxycorticosterone, 11-deoxycortisol and aldosterone. We find significant differences among these vertebrates in the transcriptional activation of their full length GRs by these steroids, indicating that there were changes in the specificity of the GR for steroids during the evolution of terrestrial vertebrates. To begin to study the role of interactions between different domains on the GR in steroid sensitivity and specificity for terrestrial GRs, we investigated transcriptional activation of truncated GRs containing their hinge domain and ligand binding domain (LBD) fused to a GAL4 DNA binding domain (GAL4 DBD). Compared to corresponding full length GRs, transcriptional activation of GAL4 DBD-GR hinge/LBD constructs required higher steroid concentrations and displayed altered steroid specificity, indicating that interactions between the hinge/LBD and other domains are important in glucocorticoid activation of these terrestrial GRs.

## Introduction

Glucocorticoids (Figure 1) regulate a variety of physiological functions including carbohydrate and protein metabolism, blood pressure, immune function and the body’s antiinflammatory processes via transcriptional activation of the glucocorticoid receptor (GR) [1–5]. The GR and other steroid receptors belong to the nuclear receptor family, a large family of transcription factors, which includes receptors for thyroid hormone, retinoids and other small lipophilic molecules [6–10]. The GR and other steroid receptors have a characteristic modular structure consisting of an N-terminal domain (NTD) (domains A and B), a central DNA-binding domain (DBD) (domain C), a hinge domain (D) and a C-terminal ligand-binding domain (LBD) (domain E) [9, 11–14] (Figure 2). The E domain alone is competent to bind steroids [11, 12, 15–18].

**Figure 1.**
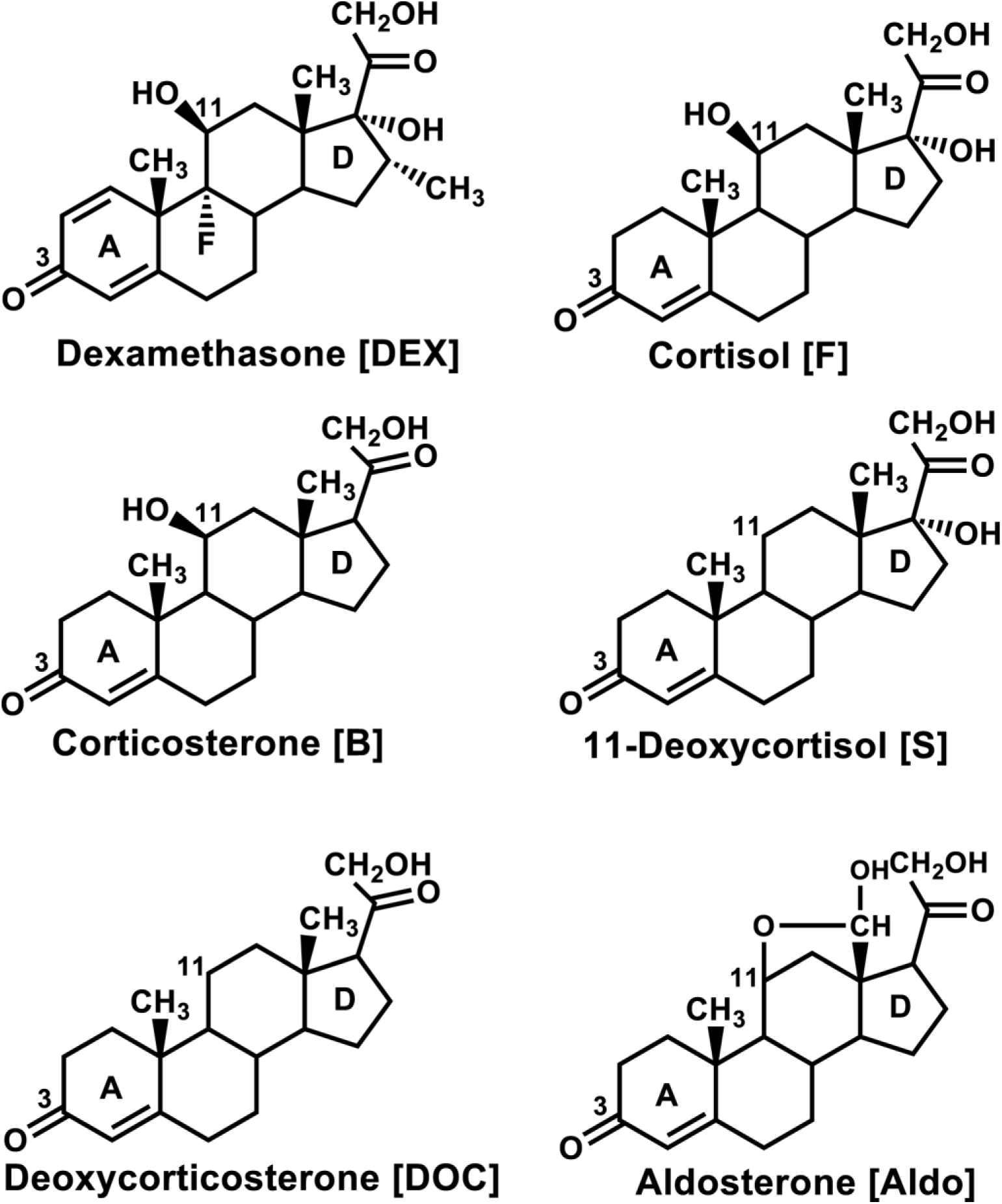
Structures of various corticosteroids. Cortisol and corticosterone are physiological glucocorticoids in terrestrial vertebrates and ray-finned fish [12, 37, 38]. Aldosterone, 11-deoxycorticosterone and 11-deoxycortisol are physiological mineralocorticoids [12, 39–41]. Aldo and DOC are weak transcriptional activators of human GR [30, 32, 36]. 11-deoxycortisol is both a mineralocorticoid and a glucocorticoid in lamprey [42].

**Figure 2.**
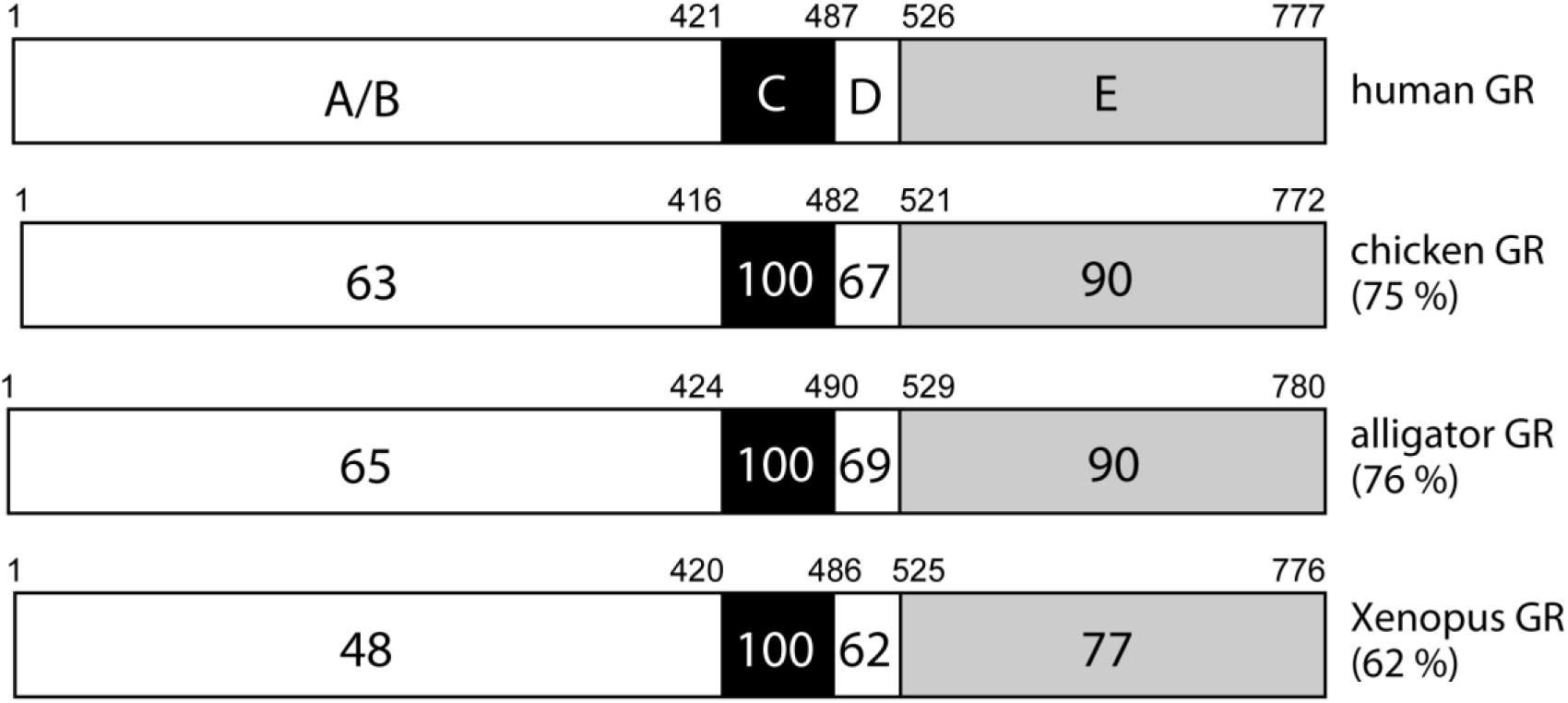
Comparison of domains in some terrestrial vertebrate GRs. GRs from human, chicken, alligator and *X.laevis* are compared. The functional A/B domain to E domains are schematically represented with the numbers of amino acid residues and the percentage of amino acid identity between the domain in the human GR and the corresponding domain in the other vertebrate GRs. For example, the entire human GR sequence is 75% identical to that of chicken GR, while domain E (LBD) on human GR is 90% identical to that of chicken GR. GenBank accession numbers: human GR (NM_000176), chicken GR (NM_001037826), alligator GR (AB701407), *X. laevis* GR (NM_001088062).

The NTD contains an activation function 1 [AF1] domain, which is a strong transcriptional activator of the GR [19–21]. Interestingly, AF1 is intrinsically disordered, unlike the DBD and LBD [21–23]. Allosteric interactions between AF1 and other domains on the GR and coactivators lead to a conformational rearrangement of AF1 that is important in transcriptional activation of the GR [23–26]. In rat GR, there is evidence that allosteric interactions between DBD and other domains regulate gene transcription [27, 28]. Recent crystal structures of the DBD-Hinge-LBD domains of other nuclear receptors [13, 22] identified allosteric signaling between the DBD and LBD domains. [27, 28].

Although dexamethasone (DEX) and cortisol (F) activation of rodent [29] and human [20, 30–33] GRs has been investigated, there has been no systematic assessment of ligand specificity among phylogenetically diverse terrestrial vertebrate GRs, such as amphibians, reptiles, birds and mammals. It is possible that more than one corticosteroid may act as a physiological glucocorticoid in terrestrial vertebrates. Reports of transcriptional activation by corticosteroids of the GR for other terrestrial vertebrates: amphibians, reptiles and birds, are limited [34, 35]. Oka et al. [34] reported half-maximal response (EC50) values for transcriptional activation of full length alligator GR by F, corticosterone (B), 11-deoxycorticosterone (DOC) and aldosterone (Aldo). The EC50s for F and B were 0.29 nM and 0.16 nM, respectively, which is consistent with the known role of these two steroids as glucocorticoids in mammals. However, the EC50s for Aldo and DOC were 2.9 nM and 2.8 nM, which is unexpected because both steroids have a lower binding affinity for human GR [30, 32]and are weak transcriptional activators of human GR [30, 36]. Similar intriguing findings for Aldo were reported for chicken GR by Proszkowiec-Weglarz and Porter [35], who found that the EC50s for transcriptional activation of chicken GR by Aldo and B were 0.8 nM and 1.8 nM, respectively, with the level of transcription due to B being about 30% higher than to Aldo. The EC50s of DOC and other corticosteroids for chicken GR and of DEX for alligator GR were not determined.

These unexpected responses of alligator and chicken GRs to Aldo and our interest in the evolution of specificity for corticosteroids in the GR in vertebrates [12, 34, 41, 43, 44] motivated us to investigate the response to a panel corticosteroids of the GR from chicken and the amphibian [*Xenopus laevis*] for comparison to human and alligator GR with the goal of clarifying the evolution of corticosteroid specificity in terrestrial vertebrates. In addition, we were interested in investigating the role of domains A-C and domains D-E [13, 21–23, 44–47] in the response of GRs to steroids. The influence of domains A-C on steroid responses for the GR has not been studied previously in non-mammalian terrestrial vertebrates. For these studies we constructed a plasmid containing the GAL4 DBD fused to the D domain and E domain of the GR (GR-LBD).

Interestingly, we found significant differences in the EC50s of these full length GRs to corticosteroids indicating that during the evolution of these terrestrial vertebrates there were changes in their response to various corticosteroids. Moreover, in the presence of corticosteroids, truncated GRs containing a GR LBD fused to a GAL4 DBD had a higher EC50 value (weaker activation) than their corresponding full length GRs, indicating altered steroid specificity among these terrestrial vertebrate GRs and that the evolution of the response of terrestrial vertebrate GRs to different steroids was complex. These differences may involve allosteric signaling between the domains D-E and other GR domains [22, 44, 45] or alterations in the binding of co-regulator proteins [24, 48, 49] or the absence of post-translational modification of domains A, B or C [49–51], or combinations of these mechanisms.

## Materials and Methods

### 2.1 Chemical reagents

DEX, F, B, aldosterone (Aldo), DOC and 11-deoxycortisol (S) were purchased from Sigma-Aldrich. For the reporter gene assays, all hormones were dissolved in dimethylsulfoxide (DMSO) and the final concentration of DMSO in the culture medium did not exceed 0.1%.

### 2.2 Construction of plasmid vectors

The full-coding regions and D/E domains of the GR from *X.laevis*, alligator, chicken and human were amplified by PCR with KOD DNA polymerase (TOYOBO Biochemicals, Osaka, Japan). The PCR products were gel-purified and ligated into pcDNA3.1 vector (*KpnI-NotI* site for human, chicken and alligator GRs, and *HindIII-NotI* site for *X. laevis* GR) (Invitrogen) for the full-coding region or pBIND vector (*MluI-NotI* site) (Promega) for D-E domains. As shown in Figure 2, the D domain begins at human GR (N487), chicken GR (N482), alligator GR (N490) and *X. laevis* GR (N486) [34].

### 2.3 Transactivation Assay and Statistical Methods

CHO-K1 cells (Chinese hamster ovary cell) were used in the reporter gene assay. Transfection and reporter assays were carried out as described previously [34, 52]. The use of CHO-K1 cells and an assay temperature of 37C does not replicate the physiological environment of *X. laevis*, alligator and chicken. Nevertheless, studies with mammalian cell lines at 37C have proven useful for other studies of transcriptional activation by corticosteroids of teleost fish GRs [53–56] and other non-mammalian GRs [35, 57, 58]. Levels of expression of the different non-mammalian GRs and their truncated counterparts may differ in CHO-K1 cells. However, comparisons of the EC50 of different corticosteroids for each GR would be valid, which is the goal of our study. All transfections were performed at least three times, employing triplicate sample points in each experiment. The values shown are mean ± SEM from three separate experiments, and dose-response data and EC50 were analyzed using GraphPad Prism. Comparisons between two groups were performed using i-test, and all multi-group comparisons were performed using one-way ANOVA followed by Bonferroni test. P<0.05 was considered statistically significant.

## Results

### 3.1 Different steroid-response for full length and truncated human, chicken, alligator and *X. laevis* GRs.

#### 3.11 Human GR

In Figures 3A and B, we show corticosteroid-inducible transcriptional activation of full length and truncated (GAL4 DBD-GR LBD) human GRs by DEX, F, B, Aldo, DOC and S. At 10^−7^ M, all corticosteroids induced transcription of full length human GR via the MMTV-reporter gene. In contrast, truncated human GR had a strong response to DEX, a much weaker response to F, a small response to B and no response to Aldo, DOC and S.

**Figure 3.**
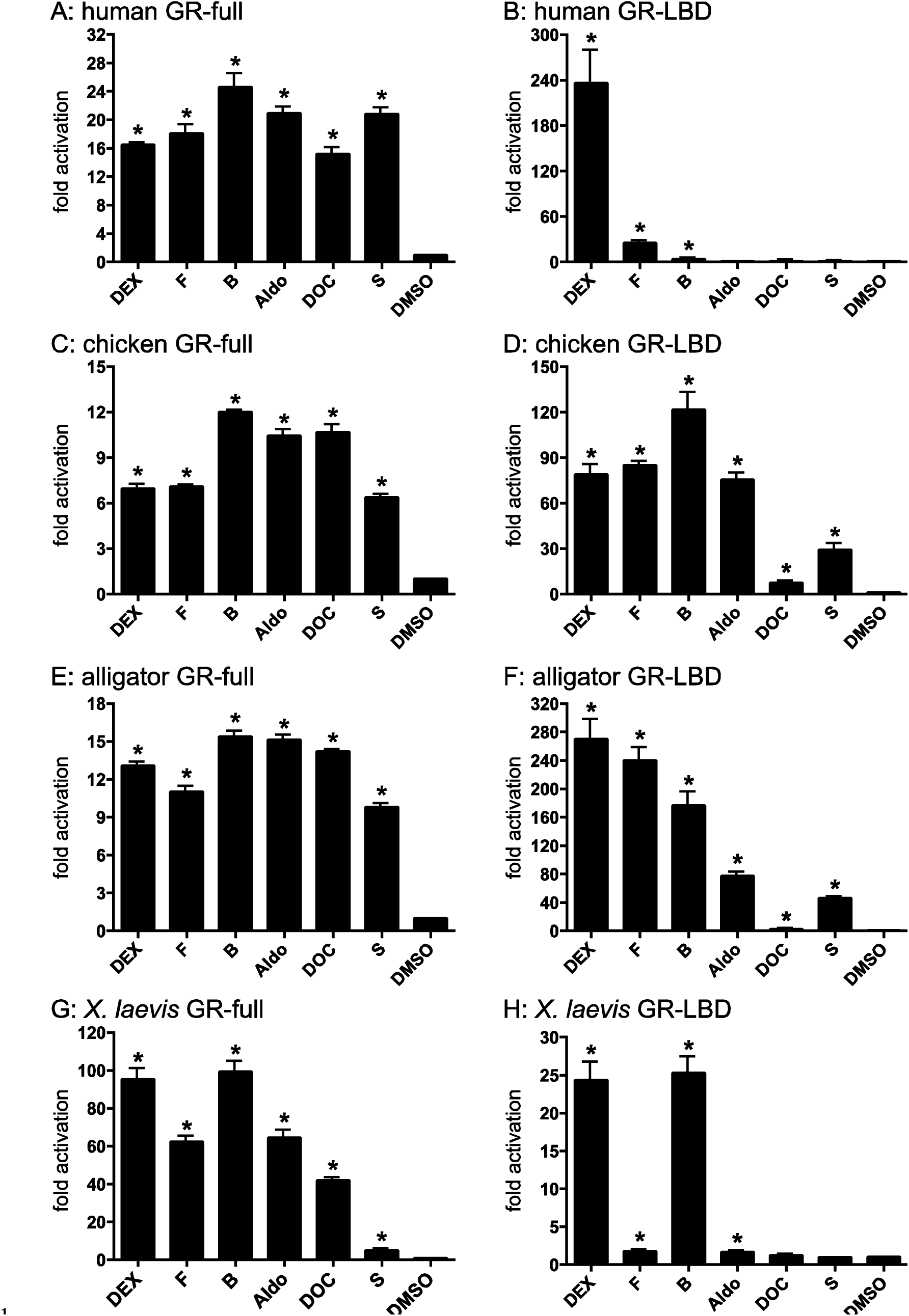
Ligand-specificities and-specificities of human, an, chicken, alligator and *X. laevis* full length GRs and LBD GRs. Full-length human GR (A), chicken GR (C), alligator GR (E), and *X. laevis* GR (G) were expressed in CHO-K1 cells with an MMTV-luciferase reporter. Plasmids for corresponding truncated GRs (human (B), chicken (D), alligator (F) and *X. laevis* (H) containing the D domain and LBD (E domain) fused to a GAL4-DBD were expressed in CHO-K1 cells with a luciferase reporter containing GAL4 binding site. Cells were treated with 10^−7^ M DEX, F, B, Aldo, DOC, S or vehicle alone (DMSO). Results are expressed as means ± SEM, n=3. Y-axis indicates fold-activation compared to the activity of control vector with vehicle (DMSO) alone as 1. Transcriptional activation of the different GRs in the presence of the DMSO control is at background level, indicating that these GRs do not have constitutive activity.

### 3.12 Chicken GR

Transcription of full length chicken GR was activated by all corticosteroids at 10^−7^ M, with a similar strong response to B, Aldo and DOC and a lesser response to DEX, F and S (Figure 3C). Truncated chicken GR was strongly activated by B, F, DEX and Aldo, with a weaker response to DOC and S (Figure 3D).

### 3.13 Alligator GR

Transcription of full length alligator GR was activated by all corticosteroids at 10^−7^ M, with a similar strong response to B, Aldo and DOC and a lower response to DEX, F and S (Figure 3E). Truncated alligator GR was strongly activated by DEX, F and B, with lower response to Aldo and S and a very weak response to DOC (Figure 3F).

### 3.14 *X laevis* GR

Transcription of full length *X. laevis* GR was activated by all corticosteroids at 10^−7^ M, with a similar strong response to DEX and B and a lower response to F, Aldo and DOC and much lower response to S (Figure 3G). Truncated *X. laevis* GR was strongly activated by DEX and B, with a much lower response to F and Aldo and a no response to DOC and S (Figure 3H).

### 3.15 EC50 values for transcriptional activation of full length human, chicken, cken, alligator and *X. laevis* GRs

Next we examined the concentration-dependence of transcriptional activation of full length terrestrial vertebrate GRs by DEX, F, B, Aldo, DOC and S (Figure 4, Table 1). Compared to the other steroids, DEX has the lowest EC50 for all of the full length GRs (Table 1). Interestingly, there are significant differences among the GRs of the EC50s for other corticosteroids, including F and B, which are the major physiological glucocorticoids in terrestrial vertebrates. For example, for full length GRs, B has a lower EC50 than F for *X. laevis* GR, while F has a lower EC50 than B for human, chicken and alligator GR.

**Figure 4.**
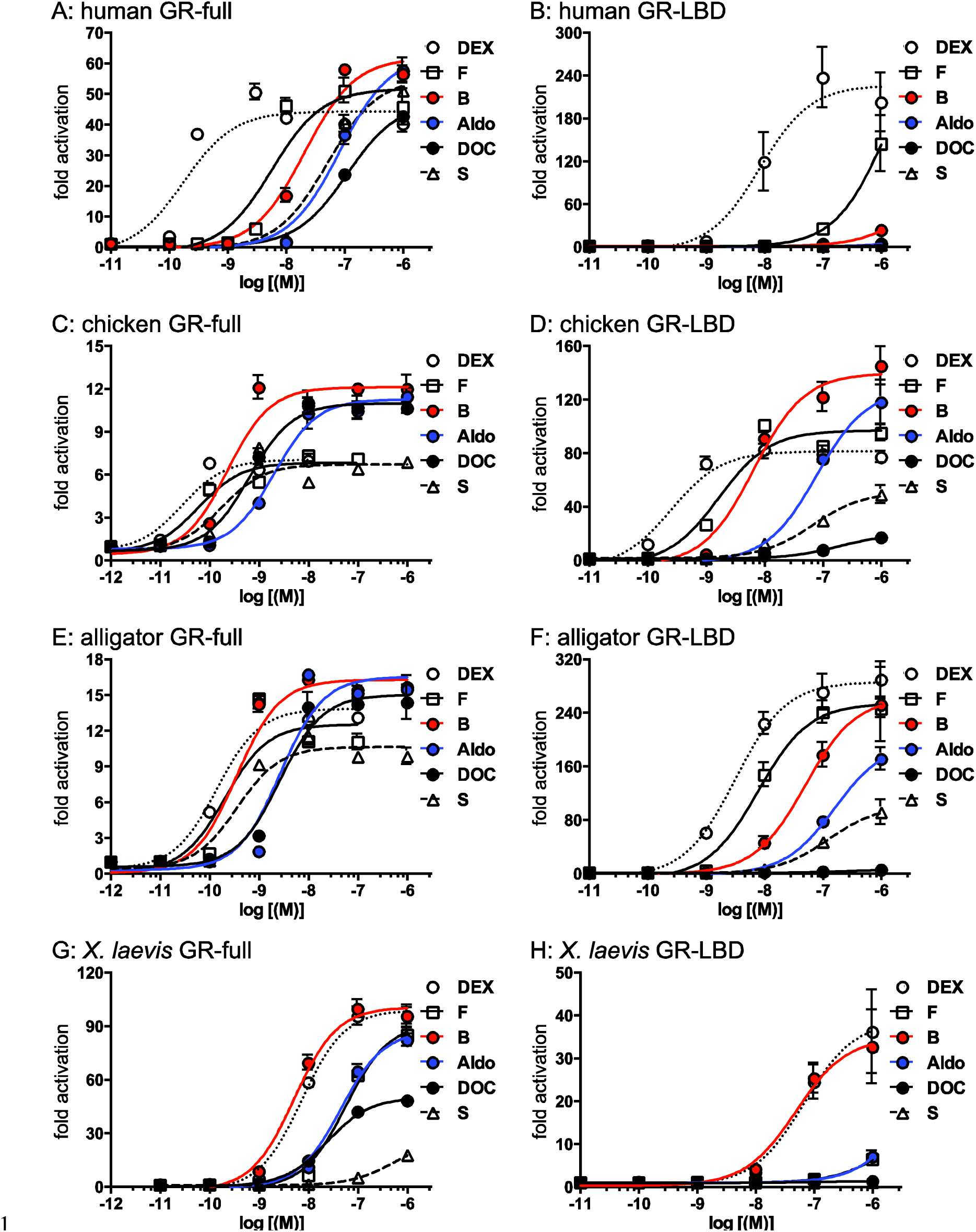
Concentration-dependent tratranscriptional activation by corticosteroids of full length and truncated human, chicken, alligator and *X. laevis* GRs. Plasmids encoding full length GRs (A: human GR, C: chicken GR, E: alligator GR, G: *Xenopus* GR) or the GAL4-DBD fused to the D domain and LBD of GRs (B: human GR, D: chicken GR, F: alligator GR, H: *Xenopus* GR) were expressed in CHO-K1 cells and treated with increasing concentrations of DEX, F, B, Aldo, DOC, S or vehicle alone (DMSO). Y-axis indicates fold-activation compared to the activity of control vector with vehicle (DMSO) alone as 1.

**Table 1.** EC50 values for transcriptional activation by corticosteroids of terrestrial vertebrate GRs.

**A. Full length GR (A-E domains)**
|  | DEX | F | B | Aldo | DOC | S |
| --- | --- | --- | --- | --- | --- | --- |
| Human GR-Full | 1.7x10<sup>-10</sup> | 5.6x10<sup>-9</sup> | 2.0x10<sup>-8</sup> | 8.2x10<sup>-8</sup> | 1.1x10<sup>-8</sup> | 5.0x10<sup>-8</sup> |
| Chicken GR-Full | 2.8x10<sup>-11</sup> | 6.0x10<sup>-11</sup> | 2.3x10<sup>-10</sup> | 2.0x10<sup>-9</sup> | 6.3x10<sup>-10</sup> | 1.7x10<sup>-10</sup> |
| Alligator GR-Full | 1.4x10<sup>-10</sup> | 2.0x10<sup>-10</sup> | 3.5x10<sup>-10</sup> | 2.7x10<sup>-9</sup> | 2.6x10<sup>-9</sup> | 3.5x10<sup>-10</sup> |
| Xenopus GR-Full | 7.3x10<sup>-9</sup> | 5.6x10<sup>-8</sup> | 5.1x10<sup>-9</sup> | 4.4x10<sup>-8</sup> | 2.3x10<sup>-8</sup> | 5.3x10<sup>-7</sup> |

**B. Truncated GR (GAL4-DBD+GR-D+E domains)**
|  | DEX | F | B | Aldo | DOC | S |
| --- | --- | --- | --- | --- | --- | --- |
| Human GR-LBD | 8.3x10<sup>-9</sup> | 1.2x10<sup>-6</sup> | - | - | - | - |
| Chicken GR-LBD | 2.5x10<sup>-10</sup> | 1.6x10<sup>-9</sup> | 6.5x10<sup>-9</sup> | 7.7x10<sup>-8</sup> | - | 6.6x10<sup>-8</sup> |
| Alligator GR-LBD | 3.1x10<sup>-9</sup> | 7.7x10<sup>-9</sup> | 4.9x10<sup>-8</sup> | 1.6x10<sup>-7</sup> | - | 1.2x10<sup>-7</sup> |
| Xenopus GR-LBD | 6.7x10<sup>-8</sup> | - | 4.8x10<sup>-8</sup> | - | - | - |

Aldo, which is a mineralocorticoid, has an EC50 of 2.7 nM and 44 nM respectively, for alligator GR and *X. laevis* GR and an EC50 of 2 nM and 82 nM, respectively, for chicken and human GR. DOC, which also is a mineralocorticoid, has an EC50 of 2.6 nM and 23 nM, respectively, for alligator GR and *X. laevis* GR, and an EC50 of 0.63 nM and 110 nM, respectively, for chicken GR and human GR. Interestingly, S has an EC50 of 0.17 nM and 0.35 nM, respectively, for chicken and alligator GR, and a much higher EC50 for human GR [50 nM] and *X. laevis* GR [530 nM].

### 3.16 EC50 values ues for transcriptional activation of truncated (GAL4 DBD-GR LBD) terrestrial vertebrate GRs

The concentration-dependence of transcriptional activation of truncated terrestrial vertebrate GRs by DEX, F, B, Aldo, DOC and S is shown in Figure 4 and Table 1. Transcriptional activation by several steroids was dramatically different among the terrestrial vertebrate GRs that lacked the A-C domains. For example, truncated human GR has a strong response to DEX (EC50 = 8.3 nM) and a very weak response to F (EC50 = 1.2 μM), and no significant response to B, Aldo, DOC or S. This contrasts to truncated chicken GR, which has nM EC50s for DEX, F and B, and a weaker but significant response to Aldo and S. Only DOC does not activate truncated chicken GR. Truncated alligator GR has nM EC50s for DEX and F, a weaker but significant response to B (EC50 = 49 nM), a weak response to Aldo (EC50 = 0.16 μM) and S (EC50 = 0.12 μM) and no response to DOC.

These results suggest that allosteric signaling between the hinge/LBD and one or more of the A, B and C domains influences the response of terrestrial vertebrate GRs to corticosteroids.

### 3.17 Analysis of a 25 residue segment on human GR and MR that influences corticosteroid specificity

Rogerson et al. [59] identified a combination of 12 amino acids in a 25 residue segment, corresponding to the c-terminus of helix 5, a β-turn and helix 6 on human GR, that could be replaced with corresponding residues from human MR to yield a hybrid GR that had an EC50 of 3 nM for Aldo. In Figure 5, we compare this segment in chicken, alligator and *X.laevis* GRs with the corresponding segments in human MR and GR. The alignment does not reveal a pattern of similarity between chicken and alligator GRs and human MR that can explain the lower EC50s that Aldo has for chicken and alligator GRs compared to human and *X. laevis* GRs.

**Figure 5.**
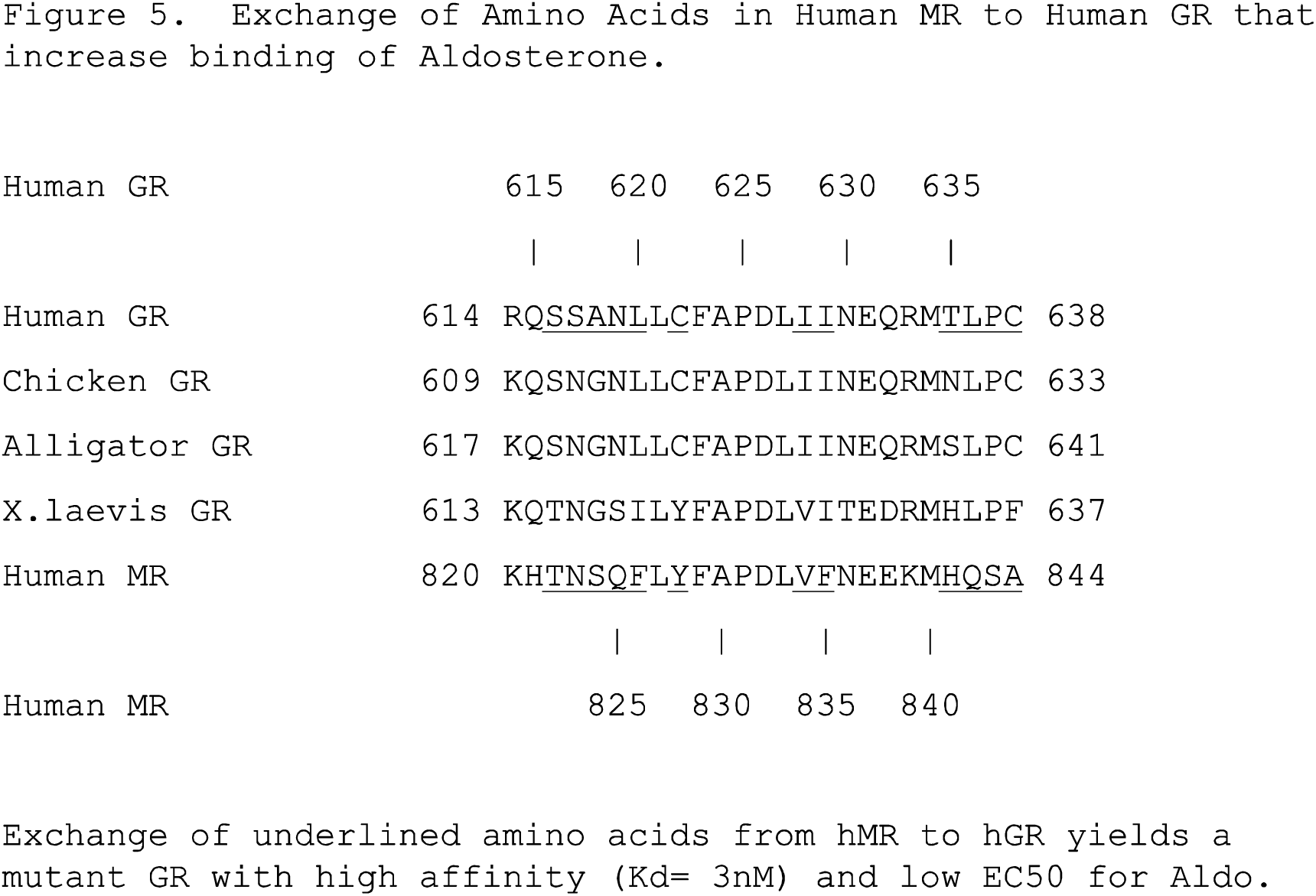
Analysis of a region in human GR and MR that is important mineralocorticoid spcificity. Rogerson et al. [59] identified a segment in human MR that could be inserted into human GR and increase its response to Aldo. We underline specific residues in the MR that replaced residues in the GR to yield an increase in the response of the GR to Aldo. Also shown is the corresponding region in chicken, alligator and *X. laevis* GRs. Residues that are conserved in all vertebrate GRs are underlined.

## Discussion

Although there are several reports of the response to different corticosteroids of the mammalian GR [20, 29–33, 36, 44], the corticosteroids that activate GRs from other terrestrial vertebrates have not been studied in depth. In birds and amphibians, B appears to be the physiological glucocorticoid [60, 61], and S has been found to be a physiological glucocorticoid in lamprey [42]. However, as discussed below, our data [Table 1] supports the presence of more than one physiological glucocorticoids in some terrestrial vertebrates. Also, our data indicate that there were changes in specificity for corticosteroids in the GR at key transitions in the evolution of terrestrial vertebrates.

As shown in Figures 3 and 4 and Table 1, we find significant differences in the response of full length GRs from *X. laevis*, alligator, chicken and humans to a panel of corticosteroids, providing evidence for the evolution of selectivity of terrestrial vertebrate GRs for F, B, Aldo, DOC and S. We confirm previous studies [34, 35] that Aldo has nM EC50s for full length chicken and alligator GR [34, 35]. This contrasts with the response to Aldo of full length human and *X. laevis* GR, for which the EC50 is 82 nM and 44 nM, respectively. The low EC50s of B for chicken GR (0.23 nM), alligator GR (0.35 nM) and *X. laevis* GR (5.1 nM) are consistent with a role for B as a physiological glucocorticoid in these vertebrates [60, 61]. We also find that DOC, another mineralocorticoid [40, 41,62], has a low EC50 for full length chicken GR (0.6 nM) and alligator GR (2.6 nM), in contrast to DOC'’s higher EC50 for human GR (110 nM) and *X. laevis* GR (23 nM). S also has a substantially lower EC50 for chicken GR (0.17 nM) and alligator GR (0.35 nM) compared to human GR (50 nM) and *X. laevis* GR (953 nM). The low EC50s of B, DOC and S for chicken and alligator GRs and of B for *X. laevis* GR leaves open the possibility that these steroids are physiological glucocorticoids in these vertebrates. There are regulatory implications for DOC and S as glucocorticoids because these steroids lack an 11β-OH group that is present in F and B [Figure 1]. Thus, DOC and S would be inert to 11β-HSD2, and could activate chicken and alligator GRs in tissues containing 11β-HSD2, which would inactivate B and F [1, 63–65].

Our studies with truncated GRs (hinge-LBD) reveal that one or more of the A, B and C domains are important in the response of terrestrial vertebrate GRs to corticosteroids. We find that compared to full length GRs, all of the truncated GRs (hinge-LBD) have substantially higher EC50s for all corticosteroids. For example, the EC50s of DEX and F for truncated human GR increased to 8.3 nM and 1.2 μM, respectively. Moreover, Aldo, B, DOC and S have an EC50 greater than 1 μM for truncated human GR. Similar changes to higher EC50s were found for the non-mammalian vertebrate GRs. Thus, F, Aldo, DOC and S have EC50s greater 1 μM for truncated *X. laevis* GR. DOC has an EC50 greater 1 μM for truncated chicken and alligator GR. Among the corticosteroids that we studied, DEX is least sensitive and DOC is most sensitive to the loss of the A, B and C domains.

There are several overlapping mechanisms that could account for stronger response to corticosteroids of full length GRs compared to that of their truncated GR counterparts. Allosteric interactions between the LBD and the NTD [44, 45, 49] or DBD [27, 47] are known to influence transcriptional activation of human and rat GRs. These allosteric interactions may be influenced by post-translational modification of the NTD by phosphorylation [49, 50] or SUMOylation [51], which also may influence binding of co-activators [24, 45, 49].

Based on Rogerson et al.’s [59] identification of a region in hMR that could be substituted into hGR and increase its response to Aldo, we analyzed the corresponding segment on chicken, alligator, *X. laevis* and human GRs for clues to differences in their responses to corticosteroids. Our analysis of this segment (Figure 5) did not find a pattern that can explain the relatively strong responses to Aldo of chicken and alligator GRs, suggesting that other mechanisms such as interactions with the LBD of the NTD on alligator and chicken GRs may contribute to the differences in their response to corticosteroids compared to human and *X. laevis* GRs.

### 4.1 Evolution

Our data indicate that there were significant changes in the response to corticosteroids during the evolution of terrestrial vertebrates. Among the species that we have studied, chicken and alligator are closest, having diverged about 150 million years ago (myr) from a common ancestor. Consistent with this close relationship, full length chicken and alligator GRs have similar EC50s for B, Aldo, S and E. In contrast, full length GR from *X. laevis*, which is the most divergent of the studied non-mammalian species, has the high EC50s for all tested corticosteroids.

It is interesting that human mineralocorticoid receptor [MR], which descended with the GR from a common ancestor [37, 66, 67], also has an interaction between domains A and B and the LBD that regulates transcriptional activation by Aldo [39, 68, 69] as does zebrafish MR [70]. The A/B domains on human and zebrafish MR can interact with each other’s LBD, indicating that this is an ancient property of the MR. This suggests that the role in transcriptional activation of the interaction between the A/B and LBD domains arose in the common ancestor of the GR and MR. Further studies of the role in transcriptional activation of the A, B and C domains on the GR and MR should provide insights into the evolution of steroid specificity in these receptors.

## Author Contributions

Yoshinao Katsu and Michael E. Baker conceived and designed the experiments and wrote the paper. Satomi Kohno and Kaori Oka carried out the research.

## Acknowledgments

We thank colleagues in our laboratories. K.O. was supported by the Japan Society for the Promotion of Science (JSPS) Research Fellowships for Young Scientists. This work was supported in part by Grants-in-Aid for Scientific Research 23570067 and 26440159 (YK) from the Ministry of Education, Culture, Sports, Science and Technology of Japan.

